# Oscillations in an Artificial Neural Network Convert Competing Inputs into a Temporal Code

**DOI:** 10.1101/2023.11.27.568876

**Authors:** Katharina Duecker, Marco Idiart, Marcel AJ van Gerven, Ole Jensen

**Affiliations:** Centre for Human Brain Health, School of Psychology, University of Birmingham Birmingham, UK; Institute of Physics, Federal University of Rio Grande do Sul Porto Alegre, Brazil; Donders Institute for Brain, Cognition and Behaviour Radboud University, Nijmegen, the Netherlands

**Keywords:** Neuronal oscillations, Multiplexing, Computer vision, Dynamical Systems

## Abstract

Deep convolutional neural networks (CNNs) resemble the hierarchically organised neural representations in the primate visual ventral stream. However, these models typically disregard the temporal dynamics experimentally observed in these areas. For instance, alpha oscillations dominate the dynamics of the human visual cortex, yet the computational relevance of oscillations is rarely considered in artificial neural networks (ANNs). We propose an ANN that embraces oscillatory dynamics with the computational purpose of converting simultaneous inputs, presented at two different locations, into a temporal code. The network was trained to classify three individually presented letters. Post-training, we added semi-realistic temporal dynamics to the hidden layer, introducing relaxation dynamics in the hidden units as well as pulsed inhibition mimicking neuronal alpha oscillations. Without these dynamics, the trained network correctly classified individual letters but produced a mixed output when presented with two letters simultaneously, elucidating a bottleneck problem. When introducing refraction and oscillatory inhibition, the output nodes corresponding to the two stimuli activated sequentially, ordered along the phase of the inhibitory oscillations. Our model provides a novel approach for implementing multiplexing in ANNs. It further produces experimentally testable predictions of how the primate visual system handles competing stimuli.

## 1 Introduction

The inclusion of convolution in artificial neural networks (ANNs) marked a significant milestone in computer vision [77]. Building on this innovation, deep convolutional neural networks (CNNs) have successfully addressed a wide range of image classification challenges, as demonstrated by Krizhevsky et al [76] in the ImageNet competition [see [114] for review]. Originally inspired by the receptive fields of neurons in visual cortex, the hierarchically organised representations emerging in these networks have been repeatedly shown to map on to those identified from human MEG and fMRI recordings of the visual ventral stream [24, 25, 42] and intracranial recordings from the non-human primate brain [73, 85, 100, 122, 121]. Despite the parallels between CNNs and the primate visual system, there are only few examples of ANNs for computer vision that have drawn inspiration from the temporal dynamics of cortical activity [31, 111]. For instance, alpha oscillations (8-12 Hz) dominate electrophysiological recordings from the human occipital lobe [1, 11], however, their computational benefit has so far not been explicitly explored in ANNs. Here, we show how embracing dynamic activations in the hidden nodes of an ANN allows the network to process competing visual inputs. This work proposes a new framework combining principles from computational neuroscience and machine learning. We show that embedding biologically inspired neural dynamics in an ANN trained for image classification, enables the network to overcome bottleneck problems when presented with multiple stimuli.

Both CNNs and the visual system of primates have a converging architecture, wherein receptive field sizes expand progressively along the hierarchy [29, 34, 37, 51, 104]. Neurons in the early layers of CNNs resemble simple cells in the primary visual cortex and possess small receptive fields that detect edges and contours in the visual input [18, 77]. The receptive fields of neurons in later layers of the CNN have more expansive receptive fields, akin to the inferior temporal (IT) cortex that has been shown to be critically involved in core object recognition [28, 39, 90, 109]. This hierarchical architecture has been argued to give rise to a bottleneck problem [16]: while early visual cortex has been shown to process visual features in parallel [22, 118, 117], object recognition has been argued have a limited capacity, which implies that the semantics of multiple visual stimuli are extracted serially [62, 97]. It has been posited that the visual and auditory systems in human and non-human primates handle simultaneously presented stimuli through “multiplexing” [3], that is, by dynamically switching between the activity patterns associated with each stimulus [19, 79] [also see[96, 95] for review]. Through this mechanism, a single neuron can contribute to the neural code of multiple stimuli [74]. This dynamic interleaving of neural responses to simultaneous stimuli requires precise temporal coordination.

One mechanism through which the neural representations of multiple objects are organised in time is phase coding, which underlies a modulation of spiking activity by ongoing low-frequency neuronal oscillations [80, 93, 103] [also see [96, 95] for review]. For example, place representations, encoded in spatially distributed firing patterns, have been shown to be ordered along the phase of hippocampal theta (4-8 Hz) oscillations[57, 56, 61, 93, 103]. Several conceptual and computational models have extended on these ideas, suggesting that the items in working memory could be segregated into cycles of ongoing gamma oscillations [81, 82, 83]. This represents a multiplexed coding scheme [83].

It has been proposed that visual perception is supported by a similar mechanism, that underlies neuronal alpha oscillations. Alpha oscillations have long been known to reflect functional inhibition [58, 71]. The strength of the inhibition waxes and wanes along the alpha cycle, such that at the peak of the oscillation, strong inhibition reduces the probability of neuronal firing in the population [44, 52]. It has been proposed that only neurons receiving sufficiently strong excitatory inputs will be able to fire at the early phases of the alpha cycle [55, 60, 59]. As the inhibition reduces toward the trough of the cycle, the neurons may fire successively according to their excitability [55, 60, 59]. In this way, the interplay between inhibition by alpha oscillations and neuronal excitability generates a temporal code.

To visualise this process, imagine the following scenario at the supermarket: While searching for the ingredients for a passionfruit martini, your gaze lands on the key element: a passionfruit, placed next to an apple (Figure 1a). With your gaze fixating on both pieces of fruit, your visual system needs to find a way to represent and process them as coherent but separate objects. As alluded to above, it has been proposed that ongoing inhibitory alpha oscillations in visual cortex serve to organise the representations of simultaneously presented stimuli competing for processing resources. It is well-documented that object-based attention is associated with increased neuronal excitability [8, 21, 66, 86, 92, 101]. Consequently, all neurons responding to the features of the passionfruit receive stronger excitatory inputs than the neurons responding to the apple and will thus overcome the alpha inhibition at an earlier phase (Figure 1b, top panel). Following the burst of activation associated with the passionfruit, refractory dynamics will momentarily deactivate its neural representation. In other words, excitation triggers inhibition, for instance, due to the activation of GABAergic [7, 72] or membrane properties such as the calcium-activated potassium current [108]. As the alpha inhibition decreases further, the neurons attuned to the apple will take over. As Figure 1b illustrates, these dynamics implement a temporal code, whereby the signals corresponding to the competing items activate successively along the alpha cycle. As the passionfruit is processed earlier in the alpha cycle, it will reach the next layer of the hierarchy with a temporal advantage over the apple. As the apple is processed in feature-selective cortex, e.g. V4, the passionfruit has already reached the object-selective cortex, e.g. IT cortex. In this way, the visual system might solve bottleneck problems through pipelining: multiple stimuli are processed in parallel, while their representations are segmented in time.

**Figure 1:**
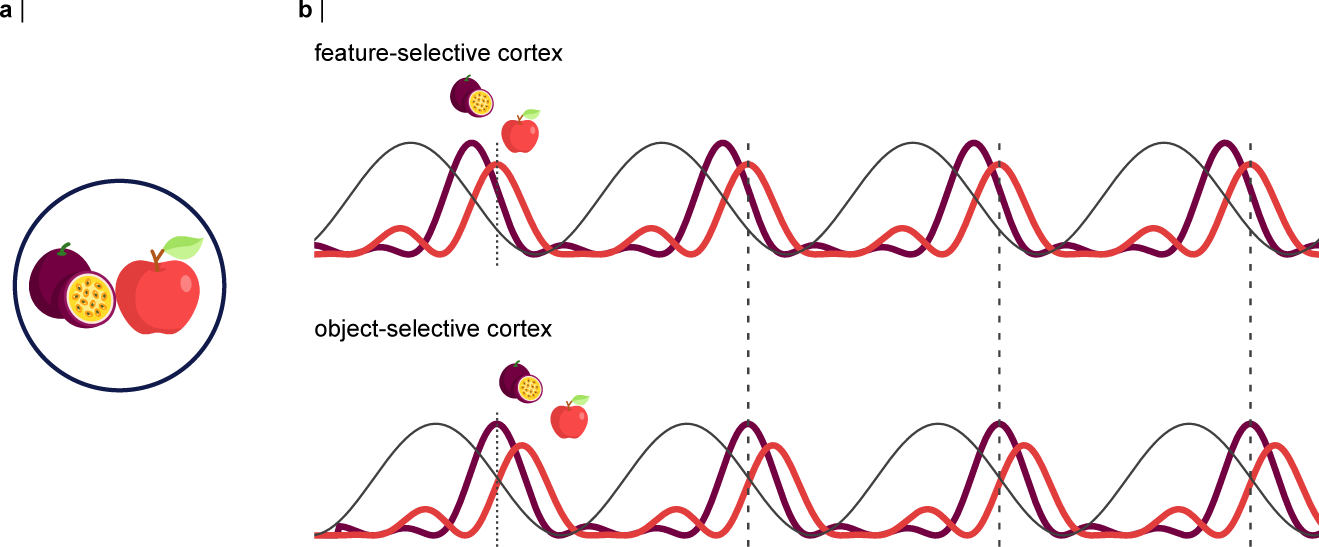
Concept: An interplay between object-based attention and neuronal alpha oscillations implements a pipelining mechanism reflected by a temporal code. **a** Example: A passionfruit and an apple are competing for the processing resources of the visual system. **b** Inhibitory alpha oscillations modulate neuronal firing rhythmically. The alpha oscillations are illustrated by the black line, the aggregate neuronal responses to the passionfruit and the apple are shown as the deep purple and red lines, respectively. As the neurons responding to the passionfruit will receive stronger excitatory inputs, they will activate at an earlier phase of the alpha cycle compared to the apple. Refraction causes a momentary inactivation of the neural representation, allowing the neurons encoding the features of the apple to activate. As the apple is processed in feature-selective cortex (e.g. V4), its representation has already been passed on to the next stage of the hierarchy, the object-selective cortex (e.g. **IT**).

**Figure 2:**
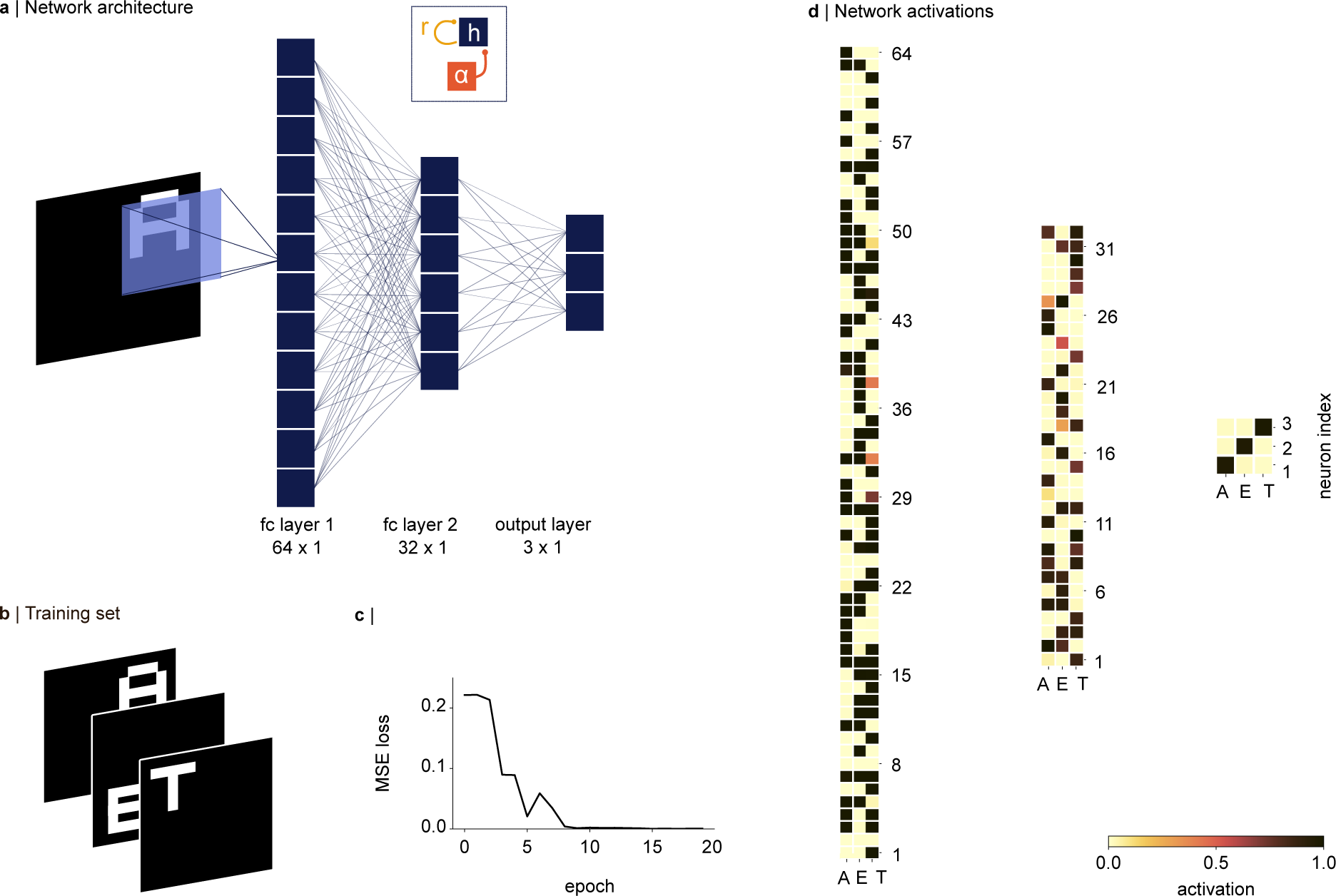
The classification problem and network architecture. **a** A network with two fully connected hidden layers was trained to classify three letters presented on a 56 *×* 56 image, in one of four quadrants (28 *×* 28). Convergence of the quadrants in the first hidden layer was implemented by sliding a 28 *×* 28 weight matrix over the image, with a stride of 28. After the training, we added a refractory term *r*, and pulses of inhibitory alpha oscillations *α*(*t*) to each hidden node *h*, as shown in the inset, and Equations 3 and 4. **b** Example inputs from the training set.**c** The network learned to classify the three letters within 10 epochs, as indicated by the mean-squared error (MSE) loss approaching 0. **d** Activations in the hidden layers and the output node in response to three inputs, presented in the three columns. The shifted sigmoid (see main text) resulted in approximately binary activations in the hidden layers. The activations in the output node demonstrate that the inputs are classified correctly.

In the following, we will present how a mechanism inspired by these concepts can be integrated into an ANN, to allow a successive read-out of competing inputs. We will refer to our model as a dynamical artificial neural network (dynamical ANN). This work serves as a proof of principle to demonstrate the basic principles and analyse the ensuing dynamics in the network. Our model relates to previous work investigating emergent dynamical and oscillatory properties in systems for information storage [50] and Recurrent Neural Networks (RNNs) for image classification [31] and sequence learning [80]. In comparison to these works, we tune the dynamics of the system such that the network is able to segment the representations of competing stimuli along the phase of ongoing oscillations in the hidden units.

## 2 Methods

To implement a dynamical ANN we first trained a two-layer network on a simple image classification task. After training, we added biologically inspired dynamics to the hidden layers motivated by alpha oscillations in the human visual system. The dynamics were not included in the training process and did not change the weights of the network.

### 2.1 Network architecture

We consider a fully connected ANN with two hidden layers, consisting of 64 and 32 hidden nodes, respectively, and an output layer with three nodes (Figure 3a). A weight matrix of size 28 *×* 28 was applied to the input (56 *×* 56) with a stride of 28, such that each node in the first layer received 4 *×* 28 *×* 28 inputs, ensuring representational invariance across the quadrants in the input. The activation *h_j_* in each unit *j* of a hidden layer was calculated as:

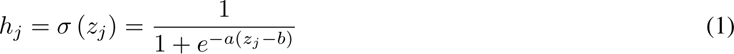

with *z_j_* the input to each hidden unit. The input *z_j_* arises from the activation in the previous layer according to *z_j_* =∑*_i_ w_ji_z_i_*, with *w_ji_* being the weight matrix connecting nodes *i* and *j*. The slope of the sigmoid was set to *a* = 2, and the sigmoid was shifted by the bias term *b* = 2.5. These parameters were fixed, such that a small input (*z ≈* 0) would result in an activation *h* close to 0, while strong inputs well above 2.5, would result in an *h* close to 1. Consequently, the activations in the hidden layers were approximately binary (“all or none”, see Figure 3d). The activation in each output node was calculated using the softmax function

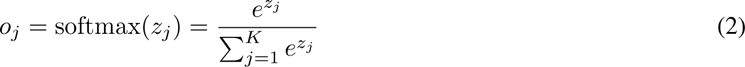

converting the inputs *z_j_* into *K* probabilities in the output layer [40]. The weights of the network were initialised according to a uniform distribution within the range [*−x, x*], where 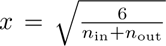 with *n*_in_ and *n*_out_ being the number of inputs and outputs to the current layer, respectively (Glorot initialization) [38]. The Adam optimiser was chosen to minimise the mean squared error loss using gradient descent [69]. The network weights were learned by backpropagating the error through the network layers (as mentioned above, the bias term was fixed at *b* = *−*2.5).

**Figure 3:**
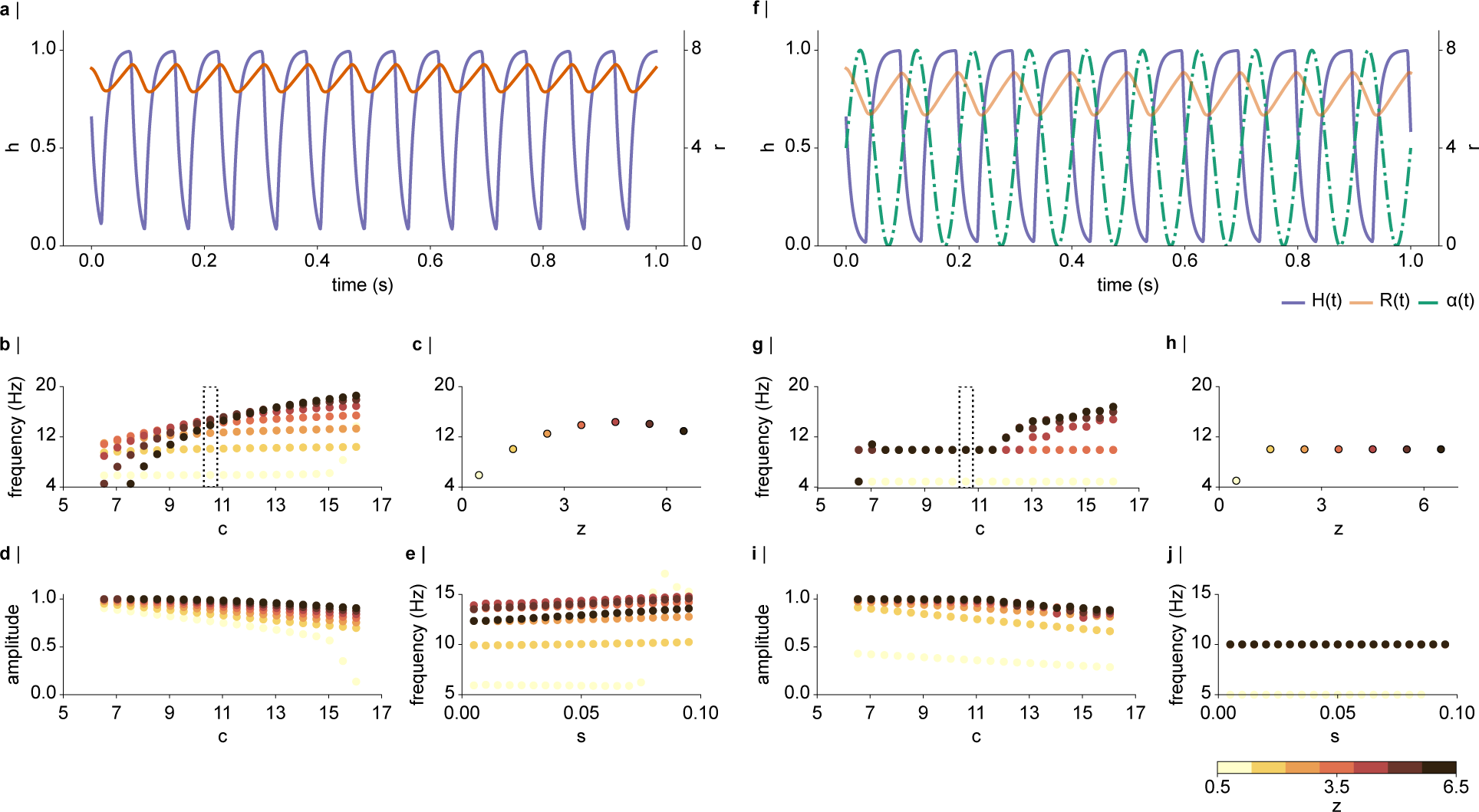
**a** Exemplary dynamics in *h* for one input *z* = 6.5, with *c* = 10 and *s* = 0.05. The interplay between *h* (purple trace) and *r* (orange trace) leads to dynamics with a period of approximately 100 ms. **b** Frequency as a function of *c* for different input values *z* (colour coded from 0.5 o 6.5). High values of *z* require larger values of *c* to generate oscillatory dynamics in *h*. **c** A value of *c* = 10 (indicated by the dotted box in **b**) leads to dynamics in the 8-12 Hz range for *z >* 0.5, whereby the frequency of the dynamics increases approximately monotonically with input size *z*. **d** Amplitude of *h* as a function of *c*, showing that for high values of *c*, *h* will not reach an activation of 1. Values of *c <* 11 seem appropriate to induce dynamics in the 10-12 Hz range that reach the sigmoid activation of 1. **e** Frequency of *h* as a function of *s*, indicating that the dynamics only change marginally for all values of *s* within a range of 0.005 to 0.1. Only for very small inputs (*z* = 0.5), a large *s* leads to fast dynamics of up to 15 Hz. **f** The dynamics of *h* and *r* shown in **a** are entrained by the 10 Hz alpha inhibition (dotted green line). **g-j** The alpha inhibition stabilises the dynamics in the 10 Hz range. **g** For large values of *c >* 11, the dynamics are able to escape the periodic inhibition and oscillate at a faster frequency. For small values of *z* = 0.5, the dynamics in *h* skip one alpha cycle, and oscillate at 5 Hz. **h** frequency as a function of *z*, for *c* = 10, showing that all nodes with inputs *z >* 0.5 are entrained by the 10 Hz rhythm. **i** The amplitude of *h* as a function of *c* in the presence of the alpha inhibition. The amplitude tends to be slightly larger than the sigmoid activation for small values of *z*. **j** In presence of alpha inhibition, *h* is robustly entrained to the 10 Hz rhythm, for all values of *s*.

### 2.2 Network dynamics in the hidden layers

We aimed to implement multiplexing by oscillatory dynamics into a fully connected ANN. To this end, after training the network on an image classification task, we added biologically inspired non-spiking dynamics to each node in the hidden layer, expressed by non-linear ordinary differential equations (ODEs). The ODEs were solved using the Euler method with a fixed time step of Δ*t* = 0.001 s. The rate of change in each hidden unit *j* was defined as:

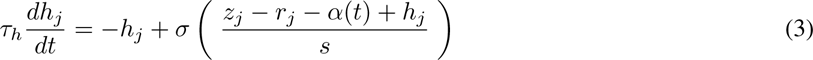

where *τ_h_* determines the timescale at which *h_j_* approaches the sigmoid activation (see Equation 1). The relaxation term *r_j_* was applied to reduce the activation of each hidden node, ensuring an intermittent activation. The dynamics of the relaxation term are given by

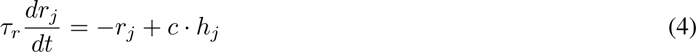

with time constant *τ_r_* defining the delay with which *r* approaches *h*. Rhythmic inhibition, mimicking inhibitory neuronal alpha oscillations in the visual system [60], was integrated into each *h_j_*, implemented as a 10 Hz sine wave

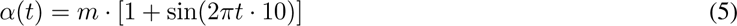

and subtracted from the input *z_j_*; with amplitude *m* being adjustable to modulate the strength and offset of the inhibition.

The selection of the parameters was informed by electrophysiological and computational constraints. For instance, the timescale of the activation *τ_h_* was set to 0.01 s, in accordance with the membrane time constant of excitatory neurons of 10-30 ms [17, 46, 113]. The time constant of the refractory term was chosen to be *τ_r_* = 0.1 s akin to afterhyperpolarization effects caused by calcium-activated potassium currents [99, 119]. The effect of parameters *c* and *s* on the dynamics will be explored below (see Figure 3).

### 2.3 Fixed points of the system

The fixed points (steady-state) of *h* and *r* are defined by setting Equations 3 and 4 to zero. Solving for *h* and *r* yields:

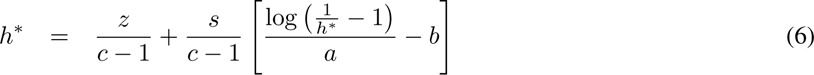

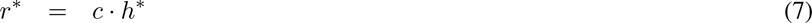

We found the fixed points numerically, using the function fsolve implemented in SciPy [112]. For small values of *s ≪* 1, the equilibrium points could be approximated as 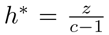 and 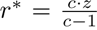. Unless otherwise specified, the dynamics of all hidden nodes in the full network were initialised at these approximated fixed points with *z* = 4 (the average input size to the first layer).

## 3 Results

The aim of this project was to show that biologically inspired dynamics will allow a neural network to handle competing inputs, despite having been trained on one stimulus at a time. To illustrate these dynamics, we trained the network to classify three letters “A”, “E”, and “T” (Figure 3a,b). The network learned to read out the images correctly within 10 training epochs (Figure 3c), with the softmax activation in the output layer approaching an activation of 1 in the node of the corresponding image (Figure 3d). Post-training, we integrated the full dynamical system described in Equation 3 and Equation 4, into the hidden layers. We will first investigate the behaviour of the individual hidden nodes.

### 3.1 Network stability and parameters

Figure 3a demonstrates how the interplay between the input *z* = 6.5 and refraction by *r* (orange trace) resulted in dynamical activations in *h* (purple trace) with a period of approximately 100 ms. As *h* increases, *r* increases with a delay. As *r* opposes *h*, this results in refraction. The adjustable parameters *c* and *s* in Equation 3 and Equation 4 were explored to identify values resulting in robust self-sustained dynamics at a frequency of about 10 Hz in the hidden nodes. Figure 3b shows how the frequency of *h* changes as a function of *c*, for *s* = 0.05, for different levels of *z* (ranging from 0.5 to 6.5 as indicated by the yellow to dark brown colour scale). The parameter *c* influences how strongly *r* grows depending on *h* (see Equation 4). Consequently, the frequency of *h* increases monotonically for increasing values of *c*. For large values of *z*, the frequency changes approximately exponentially with *c*. To induce oscillations in the 10-12 Hz range for a broad range of inputs *z*, we selected *c* = 10 (indicated by the dotted box). Figure 3c demonstrates how the frequency changes as a function of input size *z* for *c* = 10, showing that the nodes tend to oscillate at faster rhythms for larger inputs. There is a slight tendency for the frequency to plateau for very large values of *z*. This is due to *r* having to oppose a strong input while growing as a function of activation *h*, which is bounded at 1 (Equation 1). As a result, the frequency does not increase further and eventually decreases for large values of *z*.

Figure 3d demonstrates that the amplitude of *h* decreases as a function of *c*. We aimed to induce dynamics such that *h* would oscillate between 0 and the sigmoid activation of the current input, which for large values of *z* is close to 1. This amplitude was achieved for the selected *c* = 10. Note that for *z <* 4.5 the amplitude of *h* was slightly larger than the sigmoid activation of *z*. For instance, for *z* = 1.5 the activation is *h* = *σ*(*−*2 *·* (1.5 *−* 2.5)) *≈* 0.12, however, the amplitude reached values of about 0.8 (see Figure 3d). We did not find this to cause problems when integrating the dynamics into the network, as most activations in the hidden layers were outside the linear part of the sigmoid, and thus approached an activation of 1 (Figure 3c, layer 1; see below for details).

Another instrumental parameter in Equation 3 is *s*, which serves to scale the various input parameters. A small *s* will effectively increase the steepness of the sigmoid, resulting in a more step-like response. Figure 3e depicts the frequency of the node as a function of *s* and suggests that values ranging from 0.005 to 0.1 result in dynamics of about 10 Hz for a range of inputs *z*. Based on these observations, we selected *s* = 0.05.

We next explored the effects of adding pulsed inhibition at 10 Hz to the hidden dynamics (*α*(t) in Equation 3). As depicted in Figure 3f, the alpha inhibition (green dashed line) entrained the dynamics, such that *h* activated in anti-phase to the 10 Hz rhythm. Another effect of the alpha inhibition is that *r* oscillates between lower values than before. This is sensible, as the alpha inhibition and refraction *r* work together to reduce the activations in *h*. Figure 3g depicts the frequency of *h* as a function of *c*, showing that in the presence of the alpha inhibition, all hidden nodes oscillate at a frequency of 10 Hz, when 5 *< c ≤* 11, for the current input range of 0.5 *< Z ≤* 6.5. For large values of *c* (*c >* 11) the dynamics escape the alpha rhythm when *z* is sufficiently large (*>* 5) and oscillate at a faster rate. For small inputs (*z* = 0.5), *h* appears to skip one alpha cycle at a time and follows a 5 Hz rhythm. The frequency in *h* as a function of input size *z* is shown in the right panel, confirming the 10 Hz entrainment. Figure 3i shows the amplitude of *h* as a function of *c*, demonstrating that the amplitude is most stable for *c <* 11. Notably, the amplitude of *h* again reaches values above the sigmoid activation of *z* (as described above), however, we did not find this to interfere with the dynamics in the full network. Lastly, the exact value of *s* within the 0.005 to 0.1 range did not change the frequency, for all *z >* 0.5 (Figure 3j).

We conclude that the entrainment and refractory dynamics of the nodes were stable for a large range of the parameters. Based on the presented simulations, we settled on the parameters *τ_h_* = 0.01, *τ_y_* = 0.1, *c* = 10, *s* = 0.05 in all following simulations, as these produced robust dynamics at approximately 10 Hz and an activation *h* close to 1 for a wide range of inputs.

### 3.2 Alpha oscillations stabilise the dynamics in a two-layer Neural Network

After training the network on the “A-E-T” classification problem and exploring the behaviour of the individual nodes shown in Figure 3, we simulated the dynamics in the full network while keeping the weights connecting the layers fixed.

Figure 4a and b show the dynamics in the output and hidden layers of the network, in response to a single input letter, A, with *τ_h_* = 0.01, *τ_r_* = 0.1, *c* = 10, *s* = 0.05, but without any drive from the alpha oscillation (*m* = 0, Equation 5). The output node corresponding to A (blue trace) oscillates at approximately 12 Hz (Figure 4a). The leftmost panel in Figure 4b depicts the activations in *h* in layer 1 as a function of time, for *z*’s ranging from 0.5 to 6.5 (only unique values of *z* are shown). The dynamics demonstrate that the phase of the oscillations depends on the size of input *z*), leading to inconsistent phase delays between the network nodes. Due to the softmax activation introducing lateral inhibition in the output nodes, these phase delays cause a spurious intermittent activation of the output node corresponding to letters “E” and “T” (orange and green traces). The right panel in Figure 4b, depicting *h* as a function of *r*, shows the limit cycle and fixed points (indicated by the diamond-shaped scatters) for each input *z*. All units demonstrate a limit cycle behaviour but with different amplitudes.

**Figure 4:**
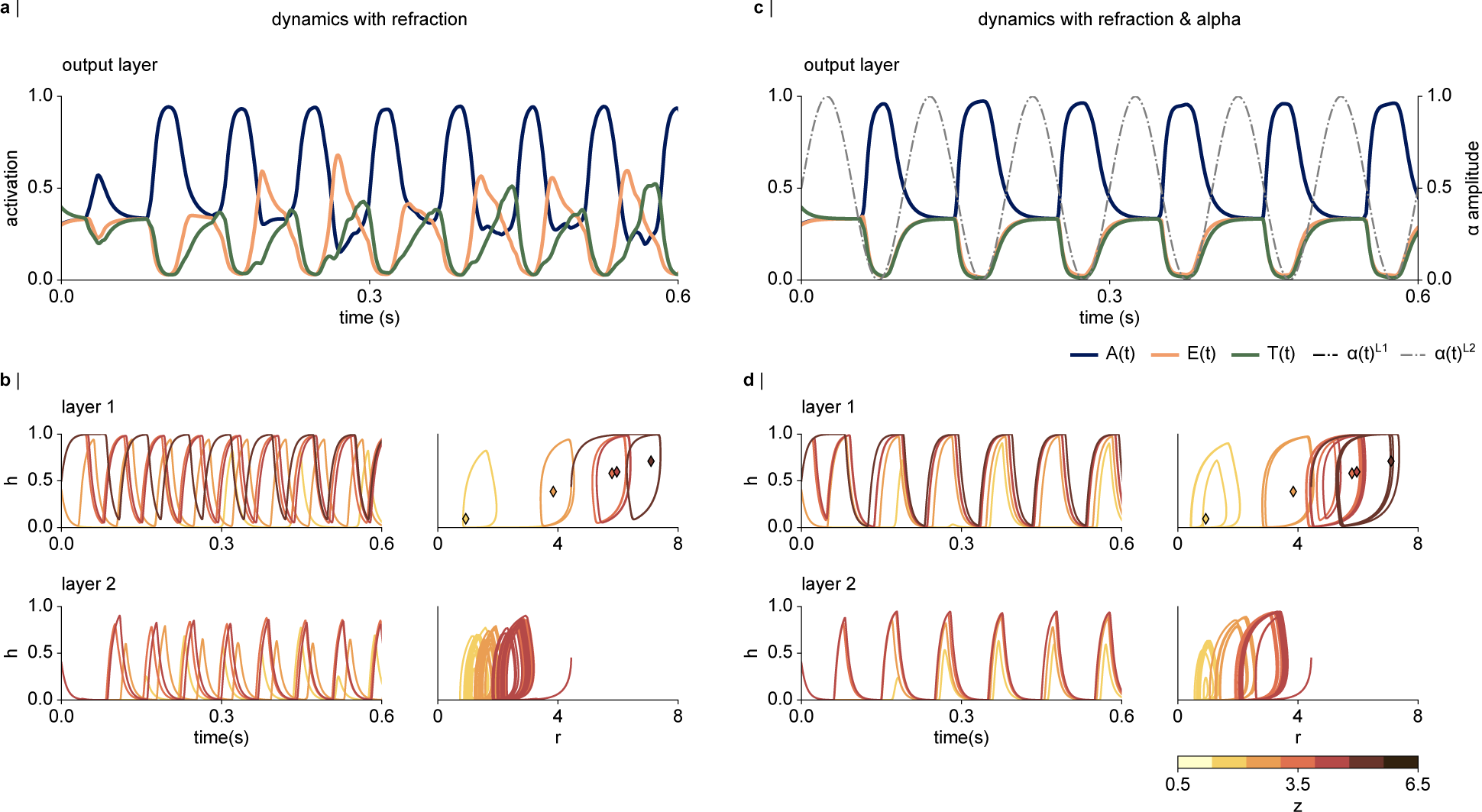
**a-b** Dynamics underlying the interplay between *h* and refraction *r* (Equation 3 and 4, without the alpha inhibition. **a** The output node corresponding to the letter A oscillates at about 12 Hz. The system spuriously activates the output node E and T. **b** (top left) The activations in the first hidden layer show that the frequency of the oscillation depends on the input size *z*, as suggested in Figure 3c. (top right) Trajectory of *h* as a function of *r*, with the fixed points indicated by the coloured diamonds. The dynamics are attracted towards a stable limit cycle, depending on the magnitudes of the input, and orbits around the fixed point. (bottom left) Activation of *h* in layer 2 as a function of time. There is a notable phase lag between the network nodes based on the input size *z*. (bottom right) The amplitude of the activation in each node appears to vary over time. **c-d** Periodic inhibition stabilises the dynamics in the system. c Periodic inhibition at 10 Hz with an amplitude of *m* = 0.5 was added to each layer. The read-out of the presented letter oscillates in anti-phase to the alpha inhibition. **d** Alpha oscillations notably stabilise the dynamics within and between the network layers, as indicated by the phase-locking between the nodes, and the stabilised H-R trajectories for the activations in layer 2.

The bottom panel shows the dynamics in the second layer. While the phase-locking between the hidden nodes in the second layer is slightly stronger compared to layer 1, activations in the nodes receiving smaller inputs appear to lag the nodes receiving larger inputs. The right-most plot, showing the relationship between *h* and *r*, again suggests a limit cycle behaviour, although the amplitude of each node appears to vary over the course of the simulation. This is likely due to the second layer receiving dynamical, non-phase-locked inputs from the first layer.

Introducing oscillatory entrainment by periodic inhibition into each layer of the network stabilises the dynamics in the entire network. Figure 4c shows the dynamics in the ouput layer, with the amplitude of the oscillatory drive set to *m* = 0.5, and a phase delay of Δ*ϕ* = 0 between the layers. Comparison to the dynamics without the oscillatory drive shows that the inhibition removes the spurious activation in the output nodes corresponding to “E” and “T” (orange and green trace). This stabilisation of the read-out underlies increased synchrony both within and between the hidden layers, as indicated in Figure 4d. In particular, the phase-locking between the activations in the first layer has been notably increased by the oscillatory drive; as shown in the time course of the activations in *h* shown in Figure 4d, top left. Comparison of the limit cycles shown in Figure 4b top right, and Figure 4d top right, demonstrates a wider limit cycle in presence of the alpha oscillations, reflecting an increased amplitude of the dynamics in *h*.

The dynamics in the second layer also show higher synchrony and stabilised trajectories *h* and *r* in presence of the alpha oscillations (Figure 4d, bottom right). This is explained by the more synchronous inputs from the first layer, and the alpha oscillations applied to the second layer. Comparison of the plots at the bottom right in Figure 4b and d reveals that the amplitude of the activation in the second layer varies less in presence of the alpha inhibition. These simulations show that the dynamics of the multi-layer network are dramatically stabilised by the oscillatory alpha inhibition.

### 3.3 Simultaneous presentation of two inputs produces a temporal code

#### 3.3.1 The bottleneck problem in absence of the dynamics

The network correctly classified individually presented stimuli after the training, as demonstrated in Figure 3c. However, when presented with two stimuli simultaneously (Figure 5a), the network produced a mixed output. The right panel in 5a shows the network activations in response to all possible inputs of stimuli. Figure 6b shows the corresponding time course to these simultaneous inputs, which was achieved using Equation 3, with *s* = 1, *r* = 0, *α*(*t*) = 0, with the network dynamics initialised at *h* = *r* = 0 in all hidden nodes. The network distributes the activations in the output layer over the nodes corresponding to the respective inputs. This suggests that the output layer produces a weighted average of the activations to both stimuli. This indicates a bottleneck problem when the network needs to classify two stimuli. In contrast, the abilities of our visual systems to recognise a stimulus do not decrease with the number of objects. As such, the visual system must be able to segment the representations of the different stimuli. As outlined above, it has been proposed that multiplexing in the visual system may underly oscillatory dynamics. In the following, we will explore how the complete dynamical network presented in Figure 4 and described by Equations 3 and 4 responds to simultaneous stimuli.

**Figure 5:**
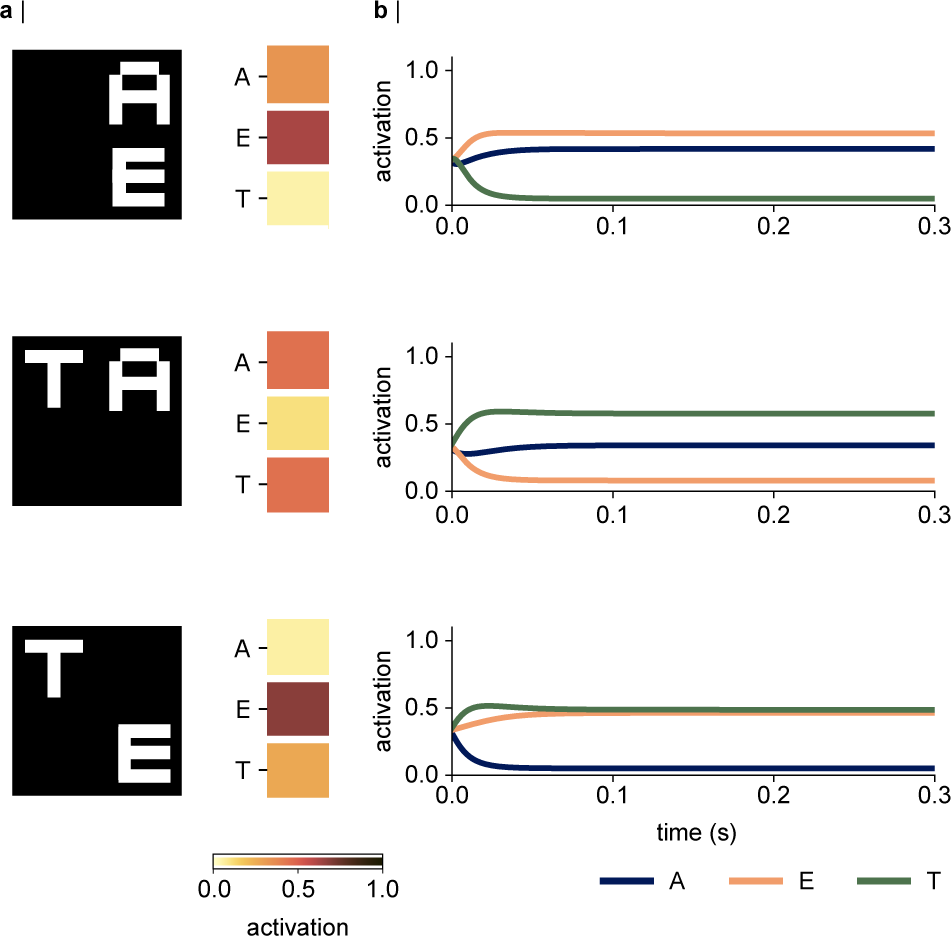
The bottleneck problem when the network receives two inputs simultaneously. **a** (left column) Exemplary input combinations of the two letters. (right) Activations in the output layer are distributed approximately evenly over the respective nodes. **b** Time course of the activation in the output layer, for *s* = 1, *r* = 0, and *α*(*t*) = 0. The shared output is a consequence of the two items competing.

**Figure 6:**
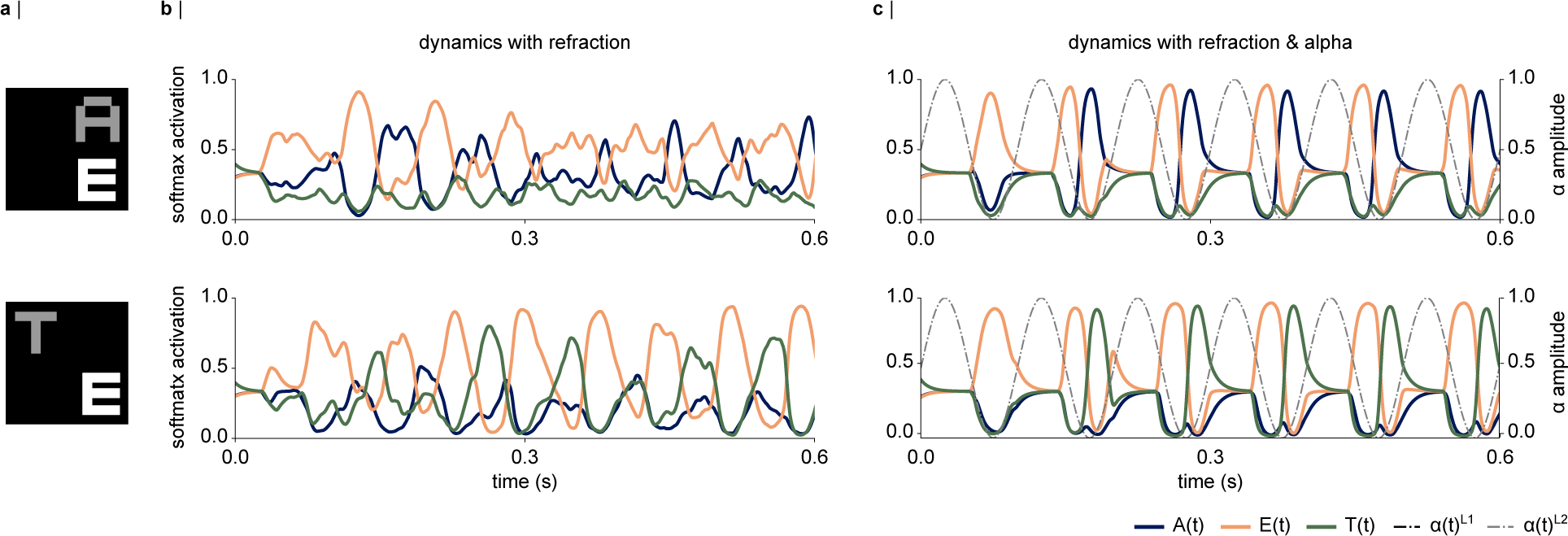
The dynamical artificial neural network (dynamical ANN) multiplexes simultaneously presented stimuli. **a** Examples of the simultaneous inputs. Attention towards the letter E was mimicked by increasing the input of all pixels belonging to E while reducing the inputs of the letters A and T. **b** The interplay between the excitation and refraction results in dynamic activations in the output layer, whereby one of the two letters is read-out at a time. However, the dynamics show notable instability, and the node corresponding to the “attended” input E activates over longer periods of time. **c** Introducing pulses of inhibition (alpha oscillations) into the network generates a temporal code in the output layer. The system first activates the node corresponding to the attended letter E. Following that, the letters are read out as a code, ordered along the phase of the alpha oscillation according to the input gain.

#### 3.3.1 Refraction and alpha inhibition allow read-out of competing stimuli as a temporal code

Figure 6b shows how the output layer of the dynamical network responds to two simultaneous inputs (“A” and “E”, and “T” and “E). To mimic spatial attention to letter E, we multiplied the pixels defining the letter E with 1.2, and the pixels defining the letters A and T with 0.8 (Figure 6a). Figure 6b shows the dynamics in the output layer, based on the simulations with parameters *τ_h_* = 0.01, *τ_r_* = 0.1, *c* = 10, *S* = 0.05, without any oscillatory drive. The node corresponding to “E” activates first and dominates the dynamics in the first 150ms (orange trace). Subsequently, the network starts to alternate between the two inputs, whereby the nodes corresponding to the presented letters reach activation levels between 0.9 and 1. As such, the interplay between activation and refraction implements a multiplexing mechanism. However, the dynamics of the multiplexed representations are unstable. For instance, in Figure 6b, top panel, the node corresponding to the letter E (orange trace) activates over longer periods of time and reaches higher activations than the node representing A (blue).

Introducing the oscillatory alpha inhibition results in stabilised dynamics, whereby the simultaneous inputs are read out as a temporal code, organised along the alpha phase (Figure 6c). Notably, the attended stimulus “E” activates first, within the first alpha cycle. In the second cycle, the system starts to generate the phase code, which further stabilises with time. We replicated these dynamics for all combinations of simultaneous inputs (two letters at a time), as shown in Figure S1.

A more detailed investigation of the multiplexing dynamics in the different layers is shown in Figure 7; for the exemplary simultaneous input E and T (as presented in Figure 6a, bottom panel). The top panels in Figure 7a and b indicate how strongly the hidden representations to the combined input correspond to the activations to the individually presented inputs. For instance, for letter “E” (orange trace) this measure of similarity was calculated as:

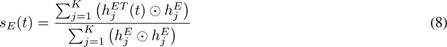

with 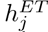 being the activation in hidden node *j* at time point *t*, to the simultaneously presented letters T and E, 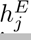 being the activation in hidden node *j* to the letter E in the trained, non-dynamical network (see Figure 3c), and *⊙* being the Hadamard operator (element-wise multiplication). This measure can be interpreted as a normalised dot product.

**Figure 7:**
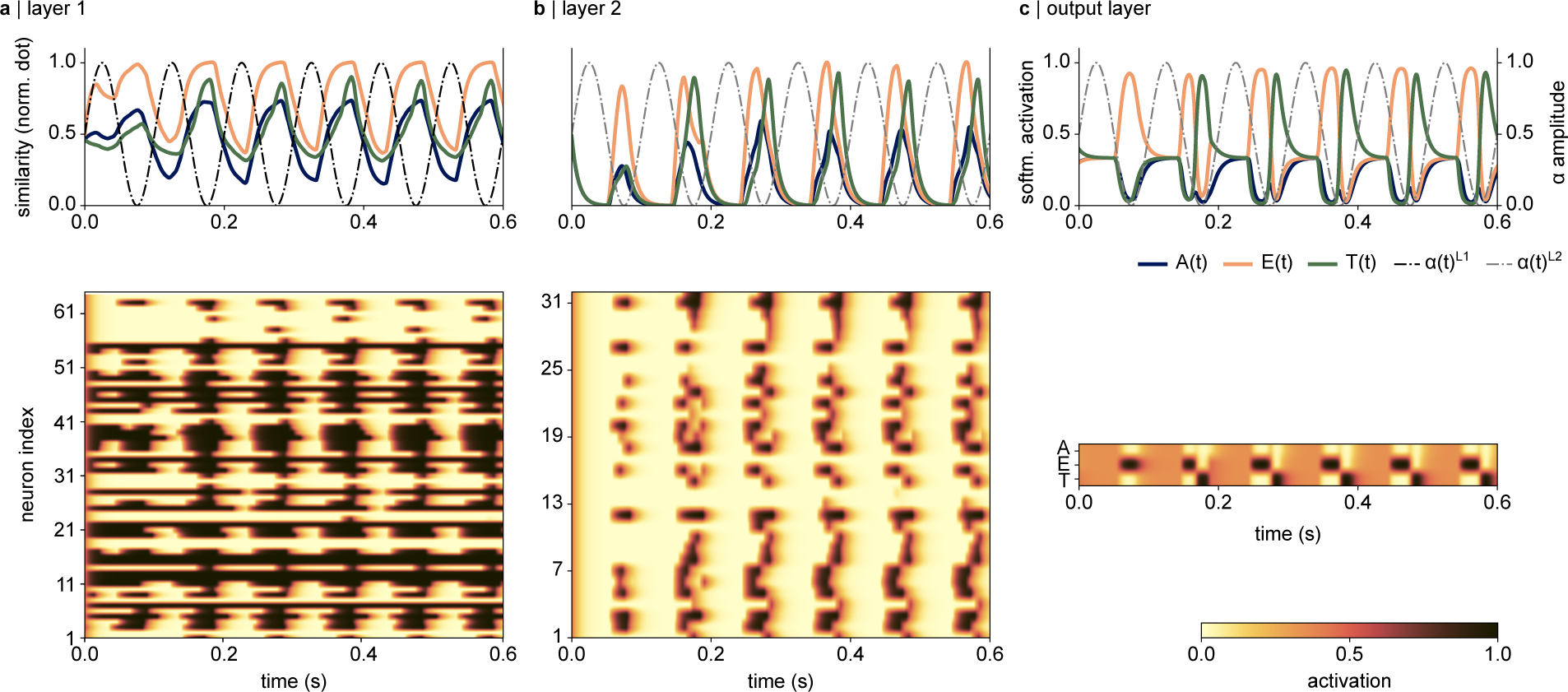
The representations of the competing inputs are segmented in the second layer of the dynamical Neural Network in response to an input image containing the letters E and T, as shown in Figure 5a, bottom. **a** (top) Layer 1: The nodes corresponding to both the letter E and the letter T activate in antiphase to the alpha inhibition, with no segregation in time. Notably, the network activates the representations to the attended letter E more strongly, and over longer periods of time. (bottom) The network activations over time suggest that a large portion of the hidden nodes activates in antiphase to the alpha inhibition. **b** Segregation of the simultaneous stimuli can be observed in the second hidden layer. (top) The normalised dot product suggests that the nodes corresponding to each input activate at different phases relative to the alpha inhibition. (bottom) The temporal segmentation observed in the top panel can also be observed as a successive activation of the nodes in the network. The alpha inhibition silences the activation in the entire layer. **c** The softmax activation and the output layer reflects a temporal code, whereby the inputs are read out along the phase of the alpha inhibition, ordered according to the magnitude of the input.

The time course of the normalized dot product indicates that the similarity between the activations in the first layer, and the representations to both letters E and T oscillates in antiphase to the alpha inhibition (Figure 7a). Notably, the similarities to both letters follow the same time course, indicating that the first layer does not segment the competing inputs. Moreover, the activations to the attended letter extend over the entire half-cycle, while the activations of the unattended letter are more short-lived. The bottom panel shows a raster plot of the magnitude of the activation in each neuron over time. The simultaneous presentation of two letters activates a large fraction of the hidden nodes in the first layer (Figure 7a, bottom). Note the rhythmic silencing of the network activations by the alpha inhibition, that affects most, but not all nodes. This implies that the first layer activates to both stimuli in parallel.

In comparison, the activations in the second layer demonstrate that the nodes responding to each letter are activated in succession: the normalised dot product between the current representations and the activations to an individual letter “E” (orange trace) precede the ones corresponding to letter T (green trace, Figure 7b). The bottom panel in Figure 7b indicates that a smaller fraction of the network is activated at each time point, and the successive activation of the hidden nodes can be observed. Finally, Figure 7c shows the read-out in the output layer, confirming that the representations of E and T are fully separated after the second cycle of the alpha inhibition (also see Figure 6c).

In sum, our simulations show how integrating dynamics driven by excitation and refraction enables a fully connected neural network to multiplex simultaneous inputs – a task it has not been trained on explicitly. This mechanism is further stabilised by pulses of inhibition, akin to alpha oscillations in the human visual system.

### 3.4 Making and breaking the temporal code: the effect of phase delay between the layers

Previous research on neural codes in the visual and auditory system has suggested that the phase of low-frequency oscillations in electrophysiological recordings carries information about the sensory input [67, 84, 91]. The communication through coherence (CTC) theory predicts that the phase relationship between two populations is critical for their communication [35, 36] (also see [2, 87] for computational implementations). While CTC was initially proposed for oscillations in the gamma-band [35, 36], related ideas have been explored for the alpha-band [14]. Based on these concepts, we next tested how different phase delays between the network layers impact the temporal code. Figure 8 shows the temporal codes for different phase delays between the alpha oscillations. Visual inspection suggests that the most robust temporal code of the two competing letters is achieved when the oscillations are synchronous in the two layers (top left), or when the second layer lags or precedes the first layer by 10 ms (top, second from left and bottom right). Notably, the temporal code breaks for larger phase delays between the layers. One might have expected an anti-phase delay between layers 1 and 2 to cut off all communication, however, the network is still able to identify at least one of the presented stimuli, albeit with high instability.

**Figure 8:**
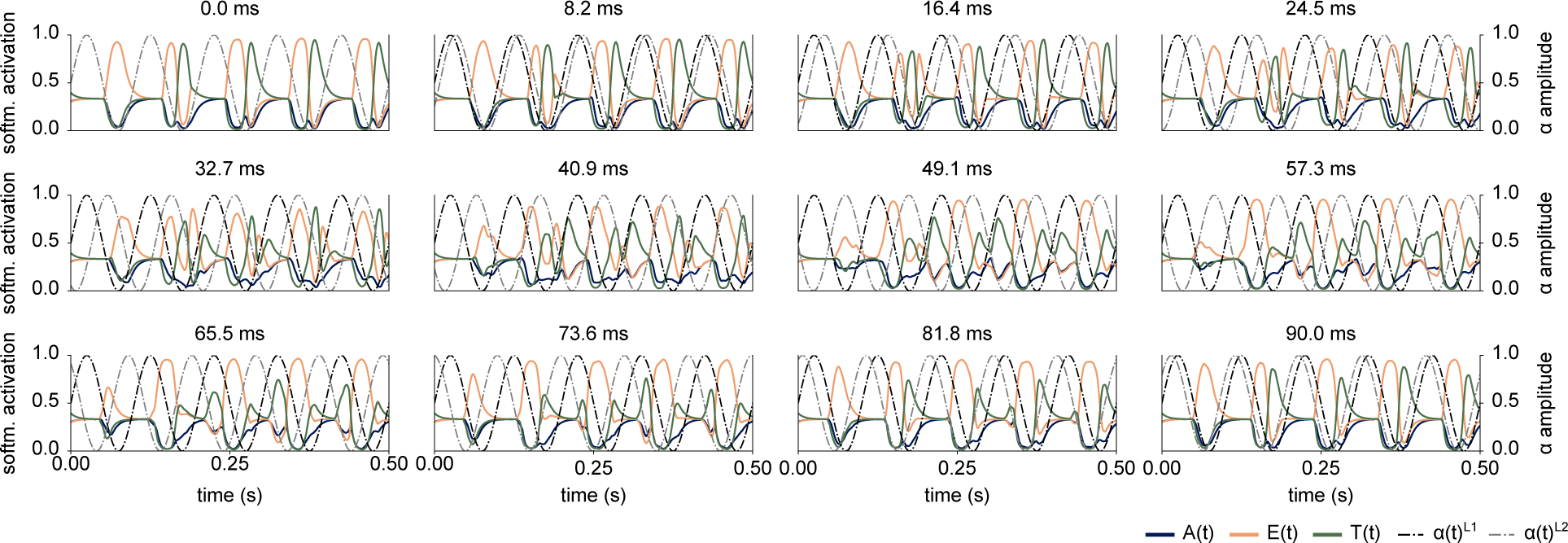
The temporal code as a function of phase delay between the layers, indicated in the title (in ms). The cleanest read-out of the two competing stimuli as a temporal code is achieved by a phase lag of zero (top left) and if the second layer precedes or lags the first layer by 10 ms.

In sum, these simulations show that the shape of the temporal code, and the number of items that can be read-out, strongly varies based on the phase delay between the layers and is most robust for synchronous oscillations in both layers.

## 4 Discussion

We here demonstrate how integrating semi-realistic neuronal dynamics into an ANN enables multiplexing during image classification. The network was first trained to classify individual letters presented in quadrants. Post-training, the network correctly recognised the individual inputs. When presented with two letters simultaneously, however, the output layer produced a mixed representation of both stimuli due to the bottleneck in the network. Adding refractory dynamics to the nodes of the network resulted in alternating activations to the simultaneously presented inputs, suggesting that the network was able to segregate the representations. However, the dynamics expressed notable instability and asynchrony within and between layers. Adding oscillatory inhibition to the layers, akin to alpha oscillations in the visual system, stabilised the dynamics in the hidden layers. When two inputs were presented simultaneously, the interplay between activation, refraction, and pulses of inhibition resulted in a stable multiplexed code, whereby the output nodes were activated sequentially based on the strength of the input.

Our simulations provide an implementation of the idea that inhibitory alpha oscillations in visual cortex serve to support the processing of simultaneously presented stimuli, by segmenting them into a temporal code [60, 59]. As the inhibition is strongest at the peak of the oscillations, only neurons receiving sufficiently strong excitatory inputs will be able to activate [55, 44, 52]. In turn, the neural representations associated with the attended stimulus will overcome the inhibition at an earlier phase of the alpha cycle than the unattended object. This allows temporal segmentation and thus multiplexing of simultaneously presented stimuli [60].

While it has been shown that simple visual features can largely be processed in parallel [22, 118, 117], object recognition has been demonstrated to be supported by serial processes [63, 97]. This indicates a bottleneck problem which has been argued to arise from the converging hierarchical structure of the visual system [16, 59, 97, 117]. By segmenting individual object representations in time, alpha oscillations have been suggested to allow the attended stimulus to pass through the network layers with a temporal advantage over the unattended stimulus (Figure 1, also see [59]). These parallel and serial properties resulting from multiplexing can be observed in our network simulation as well. In the first layer, the stimuli are represented in parallel, while the second layer begins to segment them along the phase of the oscillation (Figure 7). These simulations show how integrating biologically inspired dynamics into a neural network results in a system that can multiplex simultaneous stimuli while embracing the parallel and serial properties of visual object recognition.

Evidence for a phase-coding mechanism similar to the one presented here has been robustly observed in recordings from the rodent and human hippocampus, whereby spiking activity has been shown to be modulated along the phase of ongoing theta oscillations (4-8 Hz [61, 64, 93, 98, 103]). The order in which a sequence of inputs has been experienced, has further been proposed to be preserved in the spiking activity [61, 93] but see [80].

Intracranial recordings from the visual cortex in non-human primates have revealed that the phase of spontaneous alpha oscillations modulates spiking activity [12, 13, 30, 44] and neuronal gamma oscillations [10, 15, 88, 106]. In this context, alpha oscillations have been argued to organize visual processing in a top-down manner [68]. Intracranial and MEG recordings from the human brain have replicated the observed phase-amplitude coupling between gamma and alpha oscillations during visual processing [94, 115]. Recent human EEG recordings have additionally posited that alpha waves travelling from occipital to frontal areas are actively involved in visual processing [4, 5] (but see [125] for a critical perspective). Similarly, ECoG recordings from marmoset visual cortex have revealed travelling waves in the dorsal and ventral stream that were linked to visual performance [27] and a modulation of neural processing along the visual hierarchy following saccade initiation [65]. These reports show that alpha oscillations may propagate over cortical areas involved in visual processing to coordinate neural activity. It has so far not been determined, however, if this modulation results in successive activations of the neural representations in line with our model simulations.

Our simulations result in two testable predictions. First, we propose that neural representations activate along the phase of spontaneous alpha oscillations, ordered according to attention or salience [60, 55]. This prediction can be tested using electrophysiological recordings during visual tasks with more than one stimulus. The neural representations of each stimulus could be extracted from these data using decoding methods such as multivariate pattern analysis [49] or linear discriminant analysis [43]. For instance, using MEG, van Es et al. [32] have recently investigated the effect of ongoing alpha oscillations on the decoding accuracy of visual stimuli in a spatial attention task. The authors have demonstrated that the phase of alpha oscillations in the frontal eye field and parietal cortex of the human brain modulated the performance of the decoder. However, a phase delay between the attended and unattended stimuli has not explicitly been reported. Alternatively, these predictions could be tested based on intracranial recordings from the mouse brain. Using Neuropixels probes, spiking activity and local field potentials can be simultaneously recorded intracranially from several cortical areas in mice (and non-human primates, [53, 107]. It is well-established that the mouse visual system exhibits a hierarchical structure similar to the one observed in primates [116, 102]. As such, these data could be used to test whether spiking activity is segmented along the phase of ongoing alpha oscillations, for instance, to distinguish a figure from a background [70].

The second prediction of our model is that a stable temporal code emerges from approximately synchronous oscillations in the consecutive network layers. Recent work using concurrent iEEG and MEG recordings has suggested that interactions between alpha oscillations in prefrontal cortex and mediodorsal thalamus mediate visual performance [41]. In light of the literature on travelling alpha waves [6, 4, 5, 9, 27, 65, 124] this begs the question of whether neuronal processing along the visual hierarchy is controlled by one driving force as suggested by our simulations, e.g. the thalamus or prefrontal cortex, or a travelling wave propagating forward or backwards along the visual hierarchy. With recent advances in brain-wide recordings in mice using Neuropixels probes [53], it may be possible to investigate whether the driving force of these travelling waves can be established.

Our network relates to previous computational models that have explored the role of biologically plausible dynamics for multiplexing and inter-areal communication [3, 54, 82, 87, 110]. We expand on this work by demonstrating how multiplexing and communication through synchronous oscillatory activity can enhance the computational versatility of neural network in the context of multi-item image classification. As we aimed to provide a proof-of-principle, we trained the network on a comparably simple classification problem involving a small training set. The non-monotonic loss function in Figure 3b implies that the network might have learned to solve the problem by memorizing the inputs and could thus show low generalisability to new inputs [23]. Our aim is to expand the presented principles to CNNs with a deeper architecture that can solve benchmark image classification problems such as (E)MNIST [26, 78], CIFAR-10 [75], and ImageNet [120]. One technical detail to consider is that modern DNNs typically implement non-linearities using the rectified linear unit (ReLU) function which reduces the vanishing gradient problem in deep architectures and speeds up learning [23, 40]. Since ReLUs are not bind the activations between 0 and 1, a different set of ODEs will be needed to describe the dynamics in future versions of this model.

Alternatively, the dynamics could be integrated in additional layers with sigmoid-like activation functions, between the trained network layers, as conventionally done in spiking neural networks [105, 123]. For instance, Sörensen et al. [105] integrated spiking dynamics into a pre-trained CNN, which allowed the network to find a target stimulus in a complex natural image, an ability the model did not exhibit without the spiking dynamics. As alpha oscillations have been shown to modulate spiking activity [12, 13, 44], it would be interesting to understand to what extent oscillatory dynamics could serve to modulate activations in spiking neural networks. By incorporating spiking or non-spiking dynamics into extra layers with activations constrained between 0 and 1, the concepts presented here could be explored in pre-existing deep neural networks. In sum, future versions of this network will expand to deeper architectures and modern image classification benchmarks.

The rate of change in the network nodes was defined by a set of ODEs, following conventional practice in computational neuroscience [89]. ODEs have also found applications in the development of RNNs. For instance, in the form of neural ODEs [20], liquid-time constant neural networks [47, 48], and RNNs consisting of damped oscillators [31]. These networks had great success in learning long-range dependencies in time series data [20, 47, 48], sequences of images [80], and image classification when the pixels of the input are transformed into a time series (sequential MNIST) [31, 47]. The goal of our work, in comparison, was to convert spatially presented inputs into a time series. The dynamics in our model therefore do not provide information about the dependencies between inputs but rather serve to segment them in time. However, integrating biologically plausible dynamics into the training process would undoubtedly be interesting. Oscillatory activity in the theta, alpha, and gamma-band has been repeatedly shown to support learning and memory [for review see [33, 45]]. In line with this, Effenberger, Carvalho, Dubinin, and Singer [31] have shown that an RNN whose nodes reflect damped oscillators outperform the performance of non-oscillatory RNNs when classifying the sequential MNIST data set. For future extensions of the presented algorithm, it would be interesting to explore if and how biologically plausible dynamics could be used to support the training process.

## 5 Conclusion

We here present a proof-of-concept showing that integrating oscillatory dynamics based on excitation, refraction, and pulses of inhibition into the hidden nodes of an ANN enables multiplexing of competing stimuli, even though the network was only trained to classify individual inputs. Our simulations predict that the visual system of humans and non-human primates handles the processing of multiple stimuli by organising their neural representations along the phase of inhibitory alpha oscillations. These predictions can be experimentally tested using simple attention paradigms and electrophysiological recordings in humans, non-human primates, and rodents. Future versions of the network will include extensions to deeper architectures and modern image classification benchmarks.

## 6 Supplementary Material

**Figure S1:**
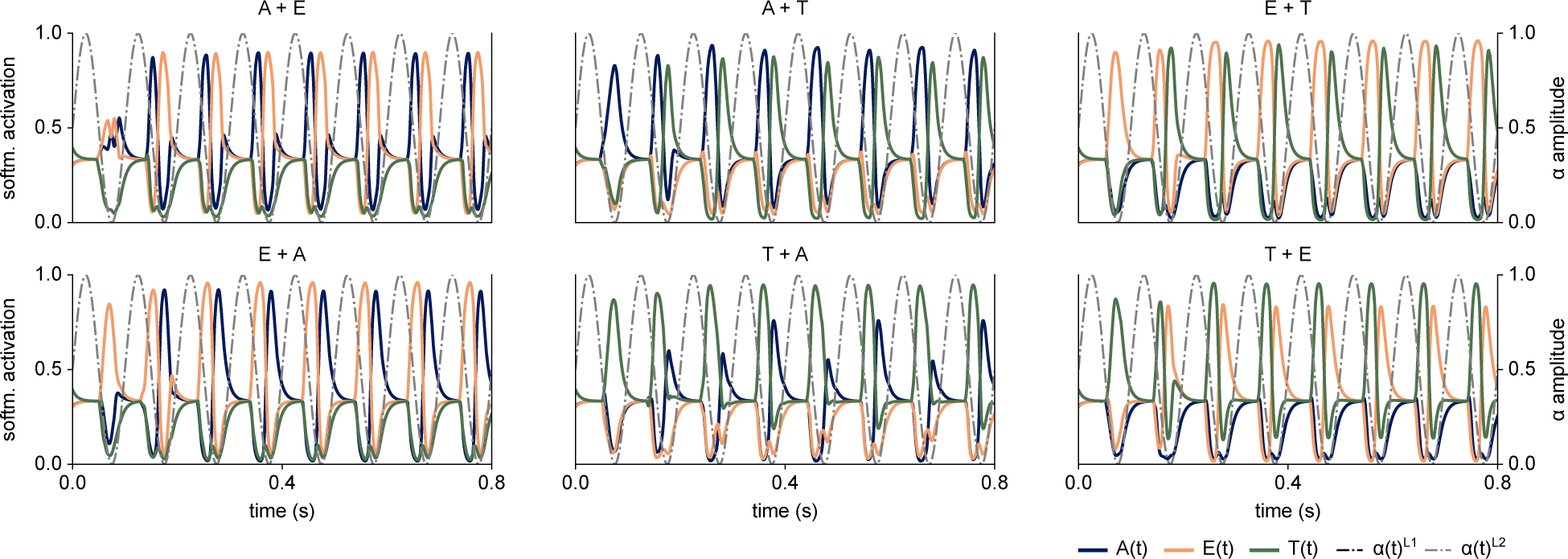
Examples of the temporal code for all input combinations. The first letter in the title is the one for which input gain has been increased.

